# Three-dimensional refractive index distributions of individual angiosperm pollen grains

**DOI:** 10.1101/353243

**Authors:** Chansuk Park, SangYun Lee, Geon Kim, SeungJun Lee, Jaehoon Lee, Taehyun Heo, Yoonjeong Park, YongKeun Park

## Abstract

Three-dimensional (3D) refractive index (RI) imaging and quantitative analyses of angiosperm pollen grains are presented. Using optical diffraction tomography, the 3D RI structures of individual angiosperm pollen grains were measured without using labeling or other preparation techniques. Various physical quantities, including volume, surface area, exine volume, and sphericity, were determined from the measured RI tomograms of pollen grains. Exine skeletons, the distinct internal structures of angiosperm pollen grains, were identified and systematically analyzed.

## 1. Introduction

Pollen grains are the male gametophytes of spermatophytes (seed plants), and they play a major role in the reproduction of various plants [1]. A pollen grain fertilizes a female gametophyte through cell-to-cell recognition and by sprouting its tube towards it. Studying various biological aspects of pollen grains has contributed to the development of multiple scientific fields. The evolution of seed plants is studied in paleontology [2], in plant ecology and reproduction [3], and in agricultural science, which focuses on plant breeding techniques [4].

Among various spermatophytes, angiosperm (flowering plant) pollen grains provide the most fruitful information regarding ecology. Angiosperms have the widest ecological niche on Earth through their active interaction with the surrounding environment during pollination. Through pollination, angiosperms influence their pollinators, including animals and insects, and even the entire ecosystem [5–7]. Therefore, the study of angiosperm pollen grains provides important clues for understanding the evolution of various species and the ecosystem.

One of the most intriguing features of angiosperm pollen grains is their morphology. The unique morphology of an angiosperm pollen grain is deeply related to its early development and survival in various environments [8–10]. There have been many attempts to study the morphology of angiosperm pollen grains. Previously, various imaging techniques were employed, including bright-field optical microscopy, scanning electron microscopy (SEM), transmission electron microscopy (TEM), fluorescence microscopy, and X-ray imaging [11–15]. However, conventional imaging techniques cannot facilitate quantitative analysis or maintaining the cell viability of a pollen grain.

Unfortunately, the abovementioned imaging techniques do not fully address three-dimensional (3D) information of pollen grains in their intact conditions. Bright-field microscopy restrictively provides low-contrast and qualitative two-dimensional (2D) imaging. TEM only provides a 2D image of a sliced sample. SEM yields 3D high resolution images of a sample, but it can only provide the surface information. Moreover, conventional techniques require either time-consuming preparation steps or complicated setups. Fixation and staining of a sample is often required for bright-field microscopy [16, 17]. Either metallic coating [11, 12] or nanometer-scale slicing [13] is required in SEM and TEM, respectively, which fundamentally limits live cell imaging using electron microscopy. Although fluorescence microscopy enables molecular-specific imaging [15, 18], the use of exogenous labeling agents compromises cell viability, resulting from phototoxicity and photodamaging [19]. Photobleaching of fluorescence proteins or dyes also prevents long-term measurements [20].

Recently, quantitative phase imaging (QPI) techniques have emerged as useful tools for label-free live cell imaging of biological samples [21, 22]. QPI exploits the refractive index (RI) of a sample as an intrinsic imaging contrast and provides unique advantages in imaging, facilitating label-free quantitative imaging of live samples [23]. Among them, optical diffraction tomography (ODT) reconstructs the 3D RI tomogram of a sample. ODT is an optical analog of X-ray computed tomography [24, 25]. In ODT, multiple 2D optical fields from a sample are measured at various illumination angles. This data is used to construct a 3D RI tomogram based on the theory of optical diffraction. 3D RI tomograms provide various quantitative information about biological samples, including cellular volume, dry mass, and protein concentration [13, 14]. ODT has been widely utilized for studying 3D structures of biological organisms, including neurons [26, 27], mammalian cells [25, 28, 29], red blood cells [30–33], eukaryotic cells [26, 34–36], immune cells [34, 37], bacteria [38–40], yeasts [41], and microalgae [42]. Recently, ODT was used to study the 3D structures of Pinus pollen grains and addressed air bag structures in grains [43]. However, label-free imaging of angiosperm has not been performed in the past.

Here we present 3D RI imaging and quantitative analyses of angiosperm pollen grains using ODT. Angiosperm pollen grains of *Lilium candidum* (lily), *Iris domestica* (leopard lily), and *Hemerocallis dumortieri* (daylily) were imaged, and their distinctive structures were analyzed systematically. Furthermore, physical quantities of the pollen grains, including volume, exine volume, surface area, and sphericity, were retrieved from the measured 3D RI distribution of the samples. This approach is label-free, requires no sample preparation, and demonstrates the potential of ODT as a tool for 3D imaging and quantitative analysis in plant biology.

## 2. Methods

A commercial ODT setup based on off-axis Mach-Zehnder interferometry (HT-1S, Tomocube Inc., Republic of Korea) was used in this study (see Figs. 1 (a) and 1 (b)). A diode-pumped solid-state laser (*λ* = 532 nm) was used as an illumination source.

**Figure 1:**
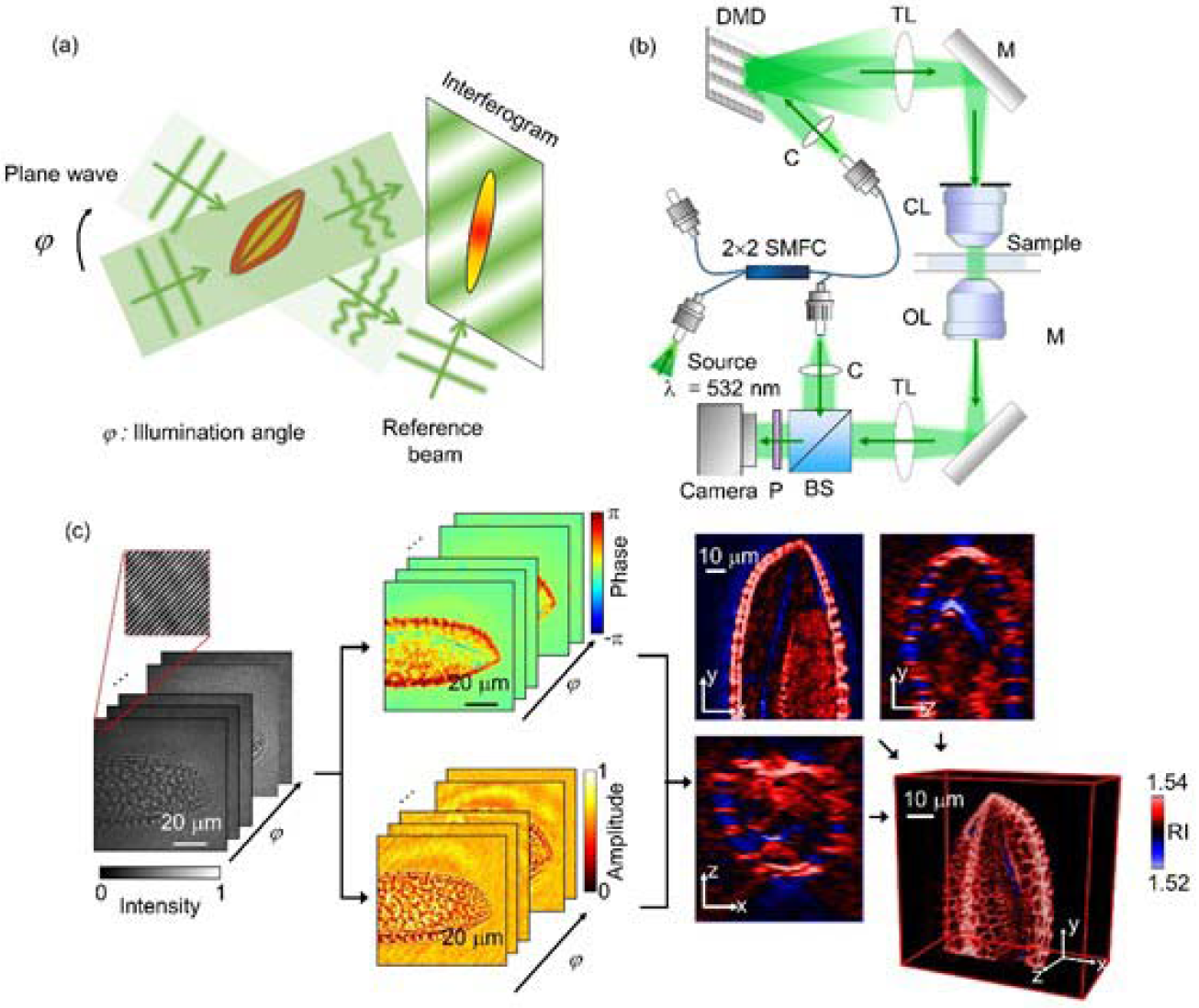
(a) Schematic diagram of the optical diffraction tomography (ODT) setup. The angle of illumination is rotated while the interferogram is recorded with the detector. (b) Optical setup for hologram measurement. 2 × 2 SMFC: single-mode fiber optic coupler, C: collimator, TL: tube lens, M: mirror, CL: condenser lens, OL: objective lens, BS: beam splitter, and P: polarizer. (c) Reconstruction procedure of the 3D RI distribution.

The beam from the illumination source was split into two arms. In a sample arm, a sample was illuminated with a plane wave, and the illumination angle was rapidly controlled using a digital micromirror device [44]. A beam scattered from a sample was projected to a camera plane with a 4-f telescopic imaging system consisting of an objective lens (60× magnification, numerical aperture = 0.8) and a tube lens (f = 175 mm). A reference arm is also projected onto a sample plane at a prescribed tilt angle. Both the sample and reference arm interfere at the image plane, and the interferogram was recorded using a CMOS camera with 150 fps frame rate (FL3-U3-13Y3M-C, FLIR Systems Inc., USA) (see Figs. 1(a) and 1(b)). In total, 49 interferograms were recorded at various illumination angles requiring 0.4 s.

From a measured hologram, the 2D optical field image, consisting of both the phase and amplitude, was retrieved by applying a field retrieval algorithm based on Fourier transform (see Fig. 1(c)) [45]. According to the Fourier diffraction theorem [24], each 2D optical field corresponding to a specific illumination angle was mapped onto a corresponding Ewald surface in 3D Fourier space [24]. Then, the 3D RI distribution of a sample was finally obtained by applying inverse Fourier transform to the filled Fourier domain. The spatial resolution of the ODT system, as calculated from the achievable spatial bandwidth, is 166 nm and 1 μm for the lateral and axial directions, respectively [46, 47]. Due to the limited numerical apertures of both the condenser and objective lens, there exists scattering signals that cannot be collected, known as missing cone information. This missing cone information was filled using an iterative regularization algorithm based on non-negativity [48]. Detailed information on ODT can be found elsewhere [25, 49].

Pollen grains from *Lilium candidum* (lily), *Iris domestica* (leopard lily), and *Hemerocallis dumortieri* (daylily) were collected from a local botanical garden (Hanbat Arboretum, Daejeon, Republic of Korea) by rubbing stamens with a cotton swab. Before imaging, pollen grains were immersed into an RI-matching oil (Series A, Cargille Labs, United States) with n = 1.53 (for *L. candidum and I. domestica*) and n = 1.54 (for *H. dumortieri*) to prevent multiple light scattering within a sample. This ensures the reconstruction of 3D RI tomography in ODT, otherwise strongly scattering samples cannot be reconstructed [50]. Since the size of a sample exceeded the field of view of the imaging system (80.4 μm x 80.4 μm), each sample was imaged twice within adjacent fields of view. Then, the 3D RI distributions obtained from different parts of a sample were manually stitched after the reconstruction procedure.

Various physical quantities of individual pollen gains can be retrieved from the reconstructed 3D RI tomogram of samples, including volume, surface area, sphericity, and exine (pollen wall) volume. The exine skeletons were distinguished from the surrounding medium by applying RI thresholds of 1.533, 1.533, and 1.527 for pollen grains from *L. candidum*, *I. domestica*, and *H. dumortieri*, respectively. The RI threshold for the exine was manually chosen such that the exine substructures were the most visible. The total region occupied by pollen grains was calculated by filling the obtained exine skeletons using a custom script using MatLab™. The exine volume *E* and the total pollen grain volume *V* were quantified by multiplying the volume of a unit voxel (4.6×10^−15^ L) and the total number of voxels occupied by the according region. The surface area of a pollen grain *SA* was calculated using a similar method. Sphericity *S* was calculated using the equation *S* = (36*πV*^2^)^1/3^/*SA*.

## 3. Results

The 3D RI distributions of pollen grains from three species of angiosperm plants were reconstructed using ODT. The 3D RI maps of the representative pollen grains for each species are shown in Fig. 2. The *x-y* cross-sectional images of the reconstructed RI tomograms are shown at various axial positions (*z* = −5, −2.5, 0, and 5 μm).

The measured RI tomograms of a pollen grain present characteristic morphological features, including exine and its substructures (tectum, columellae, and foot layer). Exine is the tough outer wall of a pollen grain and was identified as a web-like structure covering the cytoplasmic core ((i) in Fig. 2). The overall shape and size of the exine structure observed in the 3D RI tomograms are comparable to previous reports where SEM was used [9, 51–53]. Exine exhibits high RI values ranging from 1.53 to 1.54 compared to other structures. This can be explained by the fact that exine consists of sporopollenin [54], one of the most chemically stable biological polymers and a major component in the exine of plant spores and pollen grains.

**Figure 2:**
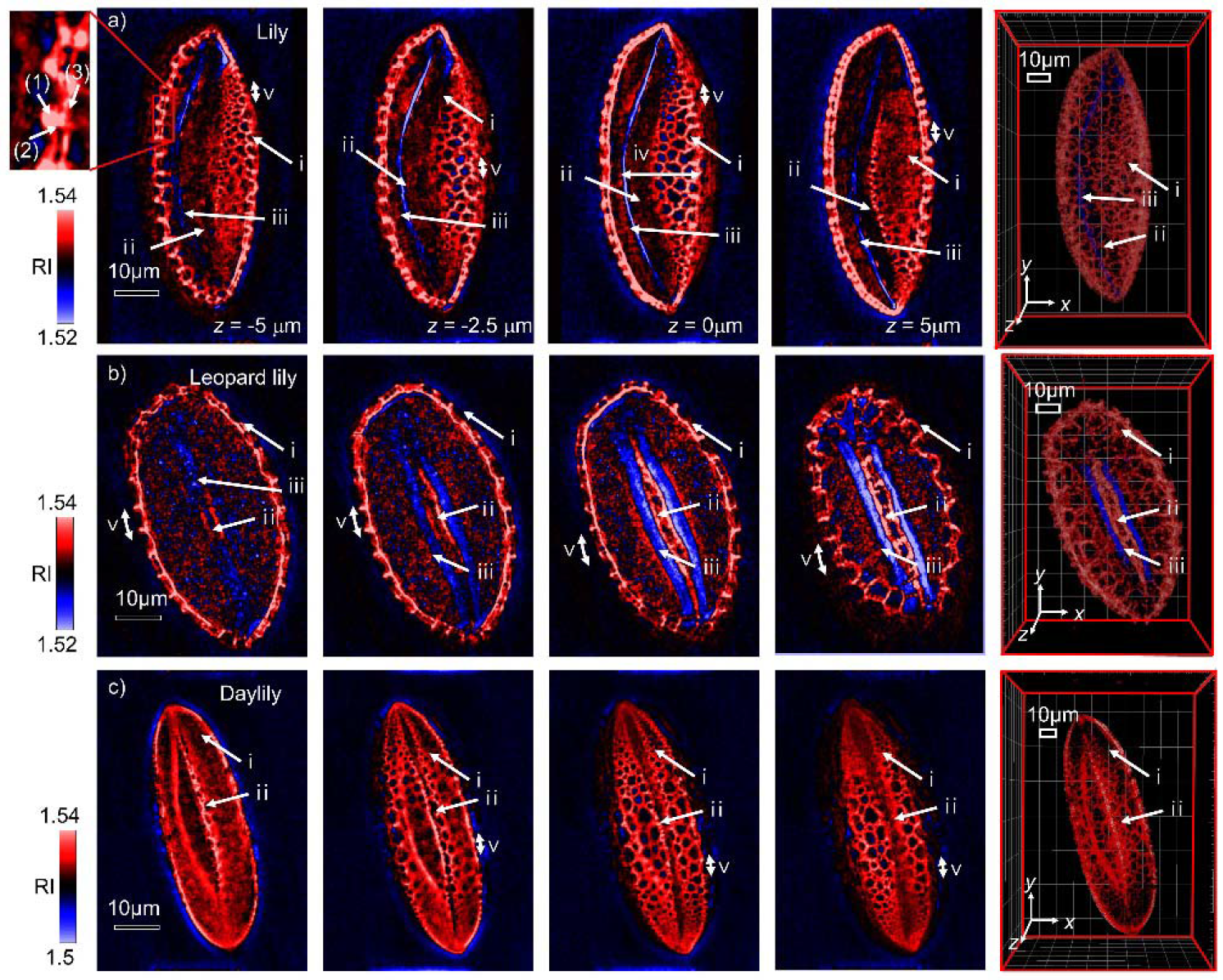
3D Volume rendering of the sample RI distribution/cross-sectional images of pollen grains from (a) *L. candidum*, (b) *I. domestica*, and (c) *H. dumortieri*. Each column includes cross-sectional images from different axial depths. The rightmost column shows a 3D volume rendering of the RI distributions. Distinctive structures of the angiosperm pollen grain are shown, including (i) exine, (ii) pollen aperture, and (iii) the lower RI region, which are assumed to be endexine and intine. Substructures of the exine were recognizable, including (1) tectum, (2) columellae, and (3) foot layer. Physical quantities of the pollen grain were measured from the cross-sectional images, including (iv) depth of the fold and (v) tectum period.

The exine substructures, including tectum, columellae, and foot layer, were also visible, as highlighted in the red box in Fig. 2(a). Tectum, the outmost layer of the exine was observed as a roof of the exine wall. The tectum was periodically distributed along the tangential direction of the pollen surface, making the surface of the pollen grain rugged. Columellae is an intermediate layer between the tectum and the foot layer and was visualized as a column connecting the tectum to the foot layer. These exine substructures are consistent with results found with SEM [55].

The pollen aperture was identified, which is the site where a pollen tube reaches out during germination ((ii) in Fig. 2) [56]. Exine wall was observed to be folded inward along the aperture. The fold is created by a process known as harmomegathy [8], which prevents dehydration of pollen grains in an arid environment. A layer with an RI of 1.517, which is lower than a surrounding medium, was visible right next to the pollen aperture ((iii) in Fig.2). These layers are assumed to be endexine and intine structures, according to the anatomical information regarding the angiosperm pollen grains from previous studies [57]. The endexine is another exine substructure located beneath the foot layer, and the intine is the inner wall of the pollen grain.

Morphological features of the pollen grains were quantified from the measured 3D RI distributions. The pollen grains from three species of angiosperms were found to be ellipsoidal. The principal axis lengths of the lily pollen grain were measured to be 66.3 μm, 46.4 μm, and 33.2 μm in descending order. The thickness of the exine wall of a lily pollen grain was 3.9 μm and was manually measured from the cross-sectional image at *z* = 0. The depth of the exine fold (35 μm) of the lily pollen grain ((iv) in Fig. 2) was directly measurable from the cross-sectional image. The *L. candidum* pollen grain had principle axes with lengths of 82.6 μm, 49.3 μm, and 41.6 μm. The exine thickness was 3.7 μm. A pollen grain from *I. domestica* showed the most distinct view of the two exine walls in contact at the pollen aperture. Moreover, a distance between the two neighboring tecta ((v) in Fig.2) of the lily pollen grain was largest (13.6 μm) among the investigated pollen grains. *H. dumortieri* pollen grain had the largest size among the three species, with principle axis lengths of 139.5 μm, 40.4 μm, and 29.9 μm. The exine thickness of the *H. dumortieri* pollen grain was 3.0 μm, which was the smallest value among the pollen grains from three species.

**Figure 3:**
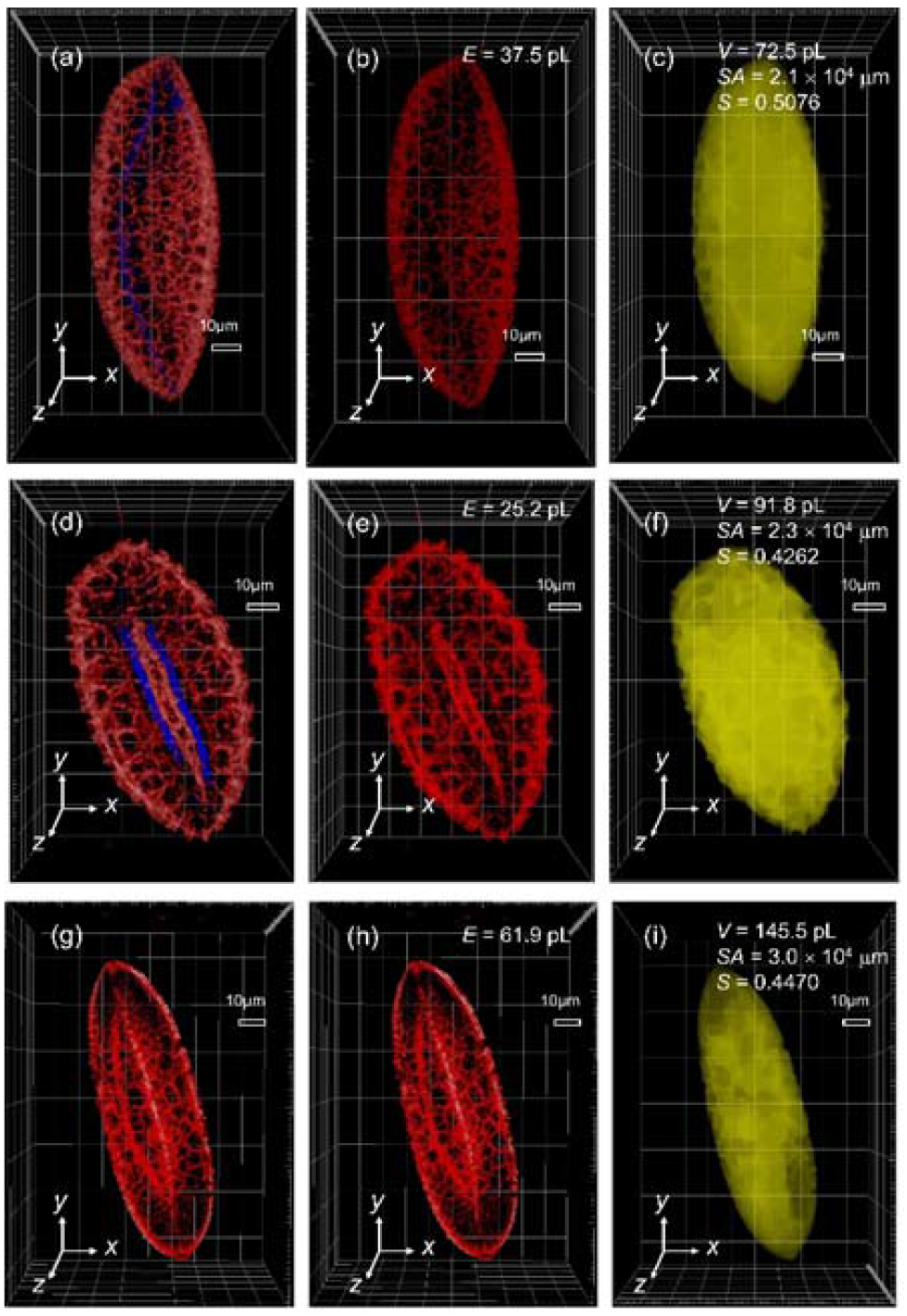
(a-c) 3D RI distribution of the pollen grains from three angiosperm species. (d-f) Exine skeletons separated by applying an appropriate RI threshold. The exine volumes E were obtained by multiplying the number of voxels occupied by the exine skeletons with the volume of a unit voxel. (g-i) Total space occupied by the pollen grains. The volumes (V) surface areas (SA) were obtained analogous to (b). Sphericity (S) of each pollen grain was calculated from V and SA.

Physical quantities including pollen grain volume *V*, surface area *SA*, sphericity *S*, and exine volume *E* were retrieved from the 3D volumetric information. Five pollens from *I. domestica*, two pollens from *L. candidum*, and one pollen from *H. dumortieri* were measured and analyzed. The representative tomograms are shown in Fig. 3. The space occupied by the exine skeleton alone and by the pollen grain was separately obtained using the RI contrast, as described in the Methods section. The results are summarized in Table 1. The volumes of the pollen grains were 86.8 ± 14.4, 91.3 ± 13.7, and 145.5 pL (mean ± standard deviation) for L. candidum, *I. domestica*, and *H. dumortieri*, respectively (Table. 1). The surface areas *S* were calculated from the isosurface of the reconstructed tomograms.

The retrieved values for the surface area were 19.9 ± 0.8, 26.3 ± 2.7, and 30.0 (× 103 μm^2^) for *L. candidum, I. domestica, and H. dumortieri*, respectively. The sphericities, calculated from the volumes and surface areas were 0.474 ± 0.034, 0.373 ± 0.030, and 0.447. The volumes of the exine structure pollen grains from different species were 31.2 ± 6.3, 28.4 ± 5.1, and 62.0 pL. The mean RI of the exine structures were 1.536, 1.536, and 1.530 for the pollen grains from *L. candidum, I. domestica, and H. dumortieri*, respectively.

**Table 1:**
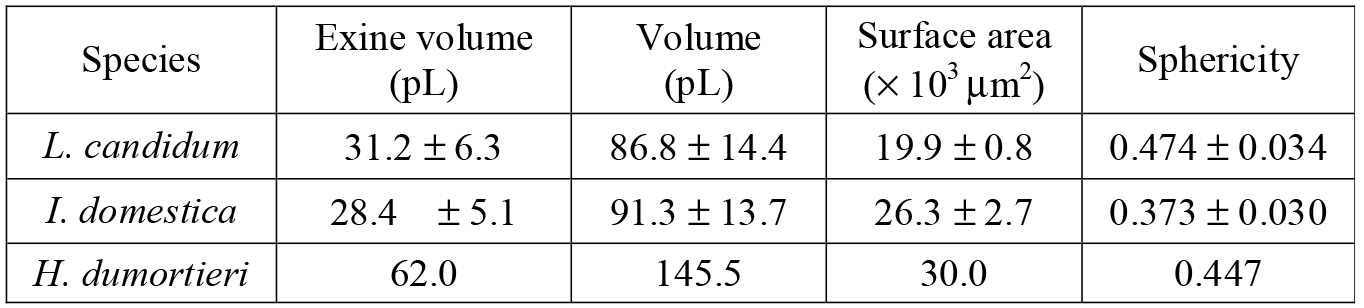
Various physical quantities obtained from the 3D RI distribution of the pollen grains from three angiosperm species.

| Species | Exine volume (pL) | Volume (pL) | Surface area ($\times 10^3 \mu\text{m}^2$) | Sphericity |
| --- | --- | --- | --- | --- |
| *L. candidum* | $31.2 \pm 6.3$ | $86.8 \pm 14.4$ | $19.9 \pm 0.8$ | $0.474 \pm 0.034$ |
| *I. domestica* | $28.4 \pm 5.1$ | $91.3 \pm 13.7$ | $26.3 \pm 2.7$ | $0.373 \pm 0.030$ |
| *H. dumortieri* | 62.0 | 145.5 | 30.0 | 0.447 |

## 4. Discussion/ Conclusion

In conclusion, label-free 3D imaging and analyses of individual angiosperm pollen grains were demonstrated using ODT. The 3D RI distribution of pollen grains provided structural information including exine and pollen aperture in pollen grains from various angiosperm plants. Furthermore, through quantitative analysis of the obtained 3D RI distribution, various physical quantities including cellular volume, surface area, and exine volume were obtained.

ODT provides 3D label-free and quantitative imaging capabilities by exploiting RI distributions in the samples. Thus, we believe ODT can offer new opportunities to investigate the diverse properties of pollen grains. For instance, ODT can provide useful insights in understanding the germination procedure of pollen grains. As is well known, the plasticity of the exine wall and the aperture of a pollen grain is linked with its ability to facilitate a volume increase of a hydrated pollen grain or germination of a pollen grain [10]. Thus, quantitative information regarding the exine wall and pollen aperture may potentially be utilized to investigate the fertilization process. Furthermore, the development of a pollen tube breaking through the pollen aperture may be directly observed using ODT without labeling agents, which has been previously investigated using fluorescence microscopy [15, 18].

Even though ODT efficiently provides 3D quantitative information on a pollen grain, ODT has limited use in molecular imaging. Therefore, subcellular organelles, such as vegetative and sperm cells could not be observed directly in ODT and requires the use of fluorescence imaging techniques [58, 59]. However, the limited molecular specificity of ODT can be overcome by various modalities in ODT, including hyperspectral [60] or polarization sensitive QPI imaging [61]. Alternatively, the correlative approach combining both ODT and fluorescence imaging [62, 63] may also provide new means to study plant biology.

